# Direction of motion decoding in mouse V1: Neuron predictive power relates to functional connectivity organization

**DOI:** 10.1101/2025.08.27.672292

**Authors:** Mario Alexios Savaglio, Christina Brozi, Eleftheria Psilou, Chryssalenia Koumpouzi, Marianna Papadostefanaki, Christos Vasilakos, Spyros Nessis, Emmanouel Smyrnakis, Vasileios Akoutas, Georgios A. Keliris, Ganna Palagina, Stelios M. Smirnakis, Maria Papadopouli

**Affiliations:** Department of Computer Science, University of Crete, Greece; Institute of Computer Science, Foundation for Research and Technology-Hellas, Greece; Archimedes Research Unit, Athena Research Center, Athens, Greece; Department of Physics, University of Ioannina, Greece; Department of Computer Science and Engineering, University of Ioannina, Greece; Brigham and Women’s Hospital and Jamaica Plain VA Hospital, Harvard Medical School, Boston, MA, USA

**Author notes:** Corresponding author: Maria Papadopouli.

**Keywords:** stimulus decoding, functional connectivity, neuronal ensembles, area V1, direction of motion

## Abstract

Variability in single neuron responses presents a challenge in establishing reliable representations of visual stimuli essential for driving behavior. To enhance accuracy, integration of responses from multiple neurons is imperative. This study leverages simultaneous recordings from a large population (tens of hundreds) of neurons, achieved through in vivo mesoscopic 2-photon calcium imaging of the primary visual cortex (V1) in mice, under visual stimulus conditions as well as in resting state (absence of stimulus). The visual stimulus consisted of 16 distinct randomly shuffled directions of motion presented to the mice. We employed mutual information to identify neurons that contain the most significant information about the stimulus direction. As expected, neurons displaying high predictive power (HPP) in stimulus decoding exhibit elevated firing event rates during stimulus presentation. Furthermore, functional connectivity among HPP neurons during visual stimulation is denser and stronger compared to functional connectivity among other visually responsive neurons. Functional connections among HPP neurons appear to form independently of distance, suggesting a distributed yet highly coordinated network. In contrast, HPP neuronal activity and functional connectivity differed significantly at resting state. Specifically, during the resting state, HPP neurons exhibited lower event rates and functional connectivity structure that was not significantly different from that of other visually responsive neurons. This suggests that HPP neurons are less susceptible to being driven simultaneously by internal brain states in the absence of a stimulus. Finally, the tuning properties of HPP neurons were unexpectedly diverse: while some were sharply tuned, others conveyed a similar amount of mutual information, despite exhibiting much weaker tuning. This study sheds light on the organization of neuronal ensembles important for decoding visual motion direction in mouse area V1, contributing to the understanding of information processing in mouse visual cortex.

## I. Introduction

Although much is known about the properties of single neuronal units, the rules by which neurons coordinate their activity to represent information about visual stimuli remain elusive. To understand why, one must consider that the responses of single units are both noisy and ambiguous: responses to the same stimulus vary considerably, and responses to different stimuli can be the same. To achieve optimal realtime performance, these ambiguities must be resolved at the level of neuronal populations via the coordinated firing of distinct neuronal ensembles. Reliable stimulus decoding in the brain is essential for guiding behavior and has been the subject of numerous studies [1]–[4]. Research in the area has recently been intensified due to the advanced imaging technologies, e.g., mesoscopic in vivo 2-photon calcium imaging, that afford exceptionally large field of view, enabling the acquisition of *simultaneous* responses across thousands of neurons [5]–[8]. Studies used decoding [5], [9], [10] to analyze how accurately visual features are represented in the brain. Information is processed in the brain by neuronal ensembles that fire synchronously, as they are likely to be more efficient at relaying information to downstream targets [11]. Recent studies demonstrate that correlations driven by the similarity in tuning of individual neurons, their response to stimuli, or higher-order correlations, affect population coding. They also play a crucial role in shaping essential functions of neural populations, such as generating codes across various timescales and aiding the transmission of information to, and interpretation by, downstream brain regions to guide behavior. A nice overview of the work on this topic can be found in [12].

Despite various efforts to uncover the underlying structures that govern reliable sensory information decoding in the brain, the *functional connectivity* properties of the *neuronal ensembles* participating in the decoding process needs to be understood better. Ensembles of neurons that fire in synchrony are likely to be more efficient at relaying shared information to downstream targets [13] and have been discussed in multiple pioneering works in relation to spontaneous or to stimulus-induced patterns of activity [3], [14]–[25]. In addition, it has been suggested that this “vocabulary space” spanned by spontaneous patterns of activity is shared with population activity patterns elicited during sensory responses [26]–[28]. This rather strong interpretation remains a matter of debate [29]. A question that drives this work is understanding how the functional connectivity between the neurons that are informative in decoding under conditions of stimulation compares to that in the resting state?

Our study leverages a rich set of data obtained in vivo using mesoscopic 2-photon calcium imaging to record essentially *simultaneously* from *thousands of pyramidal neurons* in three cortical planes, corresponding to cortical laminae 2, 3 and 4 of mouse visual cortex (Fig. 1). Here we focus on the primary visual cortex (V1). The paper employs *information-theory* to identify groups of neurons with high predictive power (i.e., **HPP** neurons) and assess how informative the neuronal activity is for decoding the visual stimulus.

**Fig. 1:**
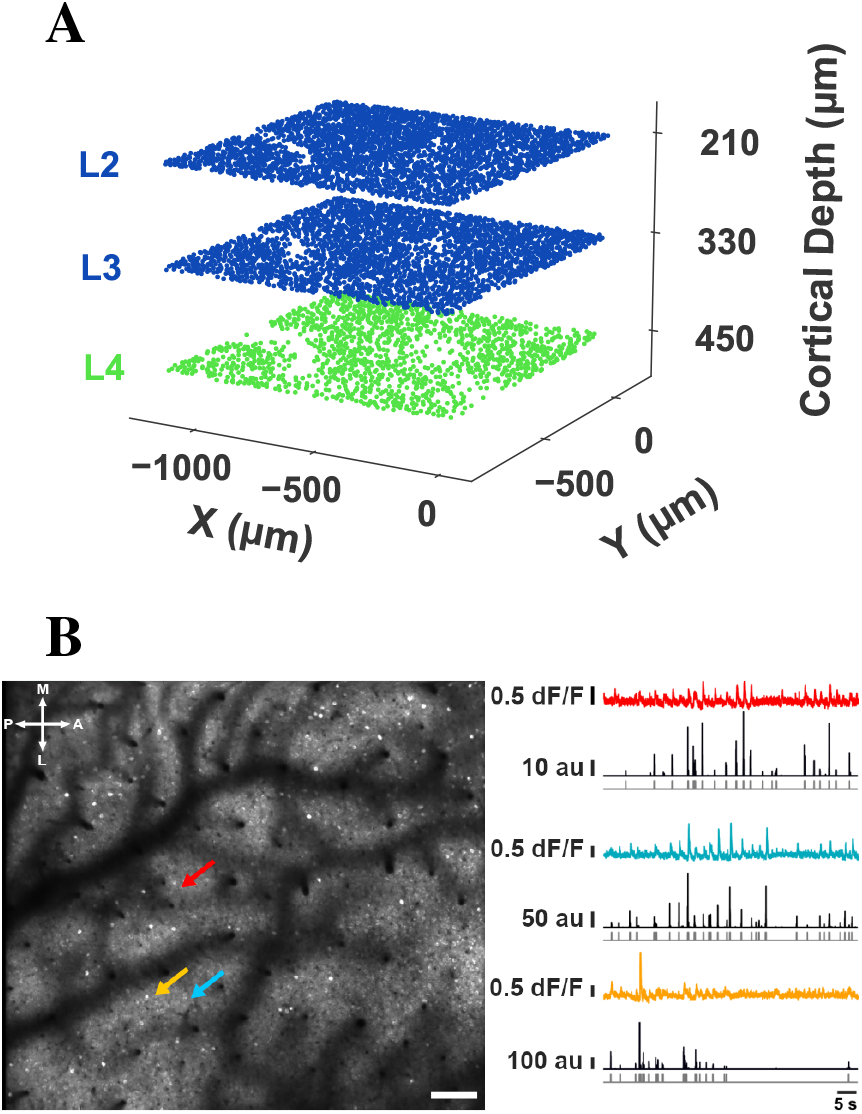
Imaging Paradigm. **A:** Illustration of L4, L2, L3 fields of view (FOVs) simultaneously acquired at 6Hz. L2/3: blue. L4: green. **B:** Example FOV acquired in L2/3 at depth 210 *µ*m. A: anterior, L: lateral, P: posterior, M: medial. Bar = 75 *µ*m. Color arrows indicate 3 example cell bodies whose traces are shown in color on the right. dF/F: fractional fluorescence change. au: the relative probability of firing in arbitrary units. Deconvolved firing probability traces shown in black below were obtained using the CaImAn algorithm [30], then thresholded to yield calcium “eventograms” that were analyzed. In what follows, we chose the threshold yielding population calcium event rates close to those reported in the literature [31], but results were robust to the choice of threshold (see suppl. Fig. 1.1 [29]). Gray traces at the bottom represent the thresholded, binarized, probability that specific imaging frames contain a calcium event (0: no event; 1: event).

We examined HPP neurons with respect to their event rate, orientation preference, and functional connectivity (e.g., degree of connectivity and length of their connections). Specifically, we used *pairwise correlations* based on STTC [32] to identify the functional connectivity, applying the methodology described in [29]. We found that most of the HPP neurons exhibit *elevated firing event rates* during visual stimulation and this pattern reverses at resting state. We found that the architecture of the HPP to HPP neuron functional connectivity under stimulation is *denser and highly distributed* compared to the architecture among the control groups, e.g., orientation-tuned neurons that are not HPP or visually-responsive but not orientation-tuned neurons. Intriguingly, during *resting state*, the functional connectivity between HPP neurons does *not* exhibit significant differences from the functional connectivity between the control populations.

The rest of this paper is structured as follows: Section II overviews the experiments, the datasets, and the neural population. In Section III, we focus on the identification of neurons with high predictive power, and profile them. Section IV analyzes the performance of decoding based on the neuronal activity profile. In Section V, we characterize the functional connectivity focusing on the HPP and the control populations. Finally, Section VI discusses our main results and future work plans.

## II. Experiments, Data Collection, And Pre-processing

This work focuses on data from the granular (L4) and supragranular (L2/3) layers in the primary visual cortex^1^ of *five adult mice*. For each mouse, we retain approximately *60-minute neuronal recordings*, during which mice were presented with stimuli videos of smoothened Gaussian noise with coherent orientation and motion (example frame in Fig. 2), consisting of waves with **16 distinct** randomly shuffled directions of motion [33]. All 16 distinct directions of motion were presented in random order in the course of a 15-sec video. Each part of this video with a *specific fixed stimulus direction* is referred to as **segment** and lasts 937.5 ms. In the following video, the 16 directions were presented in a *different* order. 240 such videos were shown *consecutively*.

**Fig. 2:**
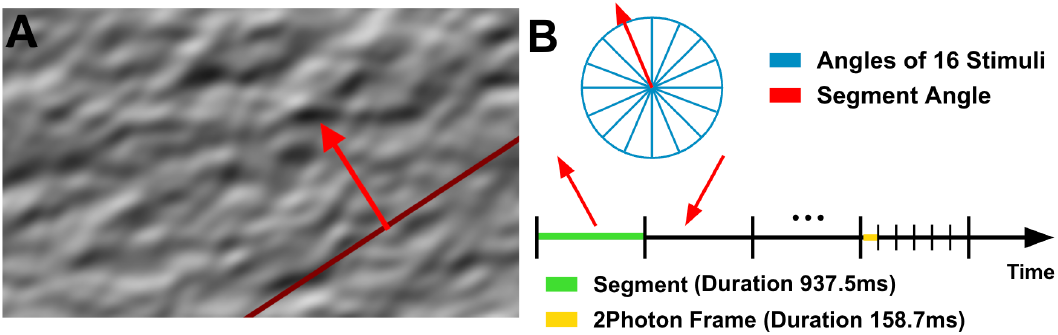
Stimulus Presentation. **A**. Example frame of “Monet” video, consisting of waves with 16 distinct randomly shuffled directions of motion, presented to the mice (i.e., stimulus). The red arrow indicates the direction of motion of the stimulus. **B**. Example of a sequence of segments, each segment with a fixed stimulus direction.

The 2-photon (2P) imaging recordings were preprocessed for motion correction and underwent automatic segmentation, deconvolution, and appropriate thresholding (see methods in Appendix) to yield calcium “*eventograms*” that were analyzed. The calcium eventogram of each neuron was subsequently used to obtain the **number of calcium event rate per segment** (*ERPS*) based on the 2P frames that coincide temporally with each Monet segment.

## III. Identification of Neurons with Information about Stimulus

Information theory has been instrumental in analyzing neurophysiological data [34] and has been used to assess the amount of information that the neuronal activity contains about a stimulus (e.g., [35]) as well as the information transmission across brain areas (e.g., [36]. To assess the amount of information about the stimulus that the firing of a neuron contains, we use the *mutual information* [34] between its *event rate per segment* and the *stimulus time series* (that indicates the angle being presented at each segment).

### Normalized Mutual Information (nMI)

The mutual information (MI) between two random variables quantifies the amount of information one variable contains about the other, i.e., it measures the *reduction in uncertainty* about one variable *when the value of another one is known*. Therefore, the mutual information *MI*(*X*; *Y*) between two jointly discrete random variables *X* and *Y*, with individual states *x* ∈ *X* and *y* ∈ *Y*, respectively, can be defined as

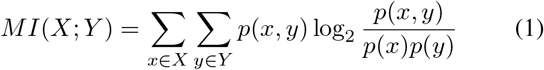

where *p*(*x, y*) is the joint probability density distribution function of *X* and *Y*, and *p*(*x*) and *p*(*y*) are the marginal probability distributions of *X* and *Y*, respectively. Normalizing the mutual information by the entropy of the stimulus facilitates a more intuitive interpretation, allowing it to be understood as the proportion of the stimulus entropy accounted for by the activity of the neuron.

Thus, the *normalized mutual information (nMI)* between two jointly discrete random variables *X* and *Y* is defined as:

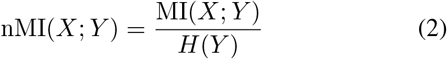

where *MI*(*X*; *Y*) is the mutual information between variables representing the event rate per segment of a neuron (*X*) and the angle of motion of the segment (*Y*), and *H*(*Y*) is the entropy of *Y*.

In our context, ERPS is treated as a discrete variable^2^. To determine the significance of the nMI between a neuron’s ERPS and the stimulus, we compare it with that obtained from a control (null) distribution. More specifically, for each neuron *i*, we randomly circularly shift its observed calcium event rate per segment time series *X*_*i*_ and then estimate the nMI with the stimulus time series *S* (i.e.,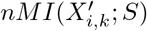), for the k-th circularly shifted instance 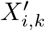 of *X*_*i*_. The above is repeated *K* = 1000 times to obtain the control values of neuron *i* (for *k* = 1, …, *K*). We then compute the *z-score* mutual information *zMI*(*X*_*i*_) of neuron *i* as:

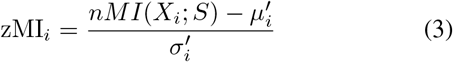

where *nMI*_*i*_ is the normalized mutual information between the observed (actual) event rate per segment *X*_*i*_ of neuron *i* with the stimulus time series 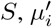 is the average normalized mutual information of the *K* control values (i.e.,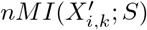), and *σ*_*i*_ is their standard deviation.^3^ The histogram of the entropy of the event rate per segment of L2/3 (L4) neurons and the mutual information with the angle of the stimulus that is presented during each segment are illustrated in Figs. 3A & 3B (4A & 4B), respectively. The control group, based on the randomly circularly shifted time series, has very low MI values with the stimulus, e.g., mean 0.012 for L2/3 neurons. A *“theoretical”* neuron with exclusive directional selectivity, which fires at all frames during segments when the direction of the stimulus is its “preferred angle”, while it remains silent at all frames of all other segments (~0.40 Hz) exhibits an MI equal to ~0.34. An example (observed) orientation-tuned neuron with a similar mean event rate (~0.39 Hz) has an MI of ~0.16. The mutual information of the L2/3 (L4) neurons with the stimulus has a skewed distribution with large tails, with a mean of 0.061 (0.062), respectively (Fig. 3B & 4B).

**Fig. 3:**
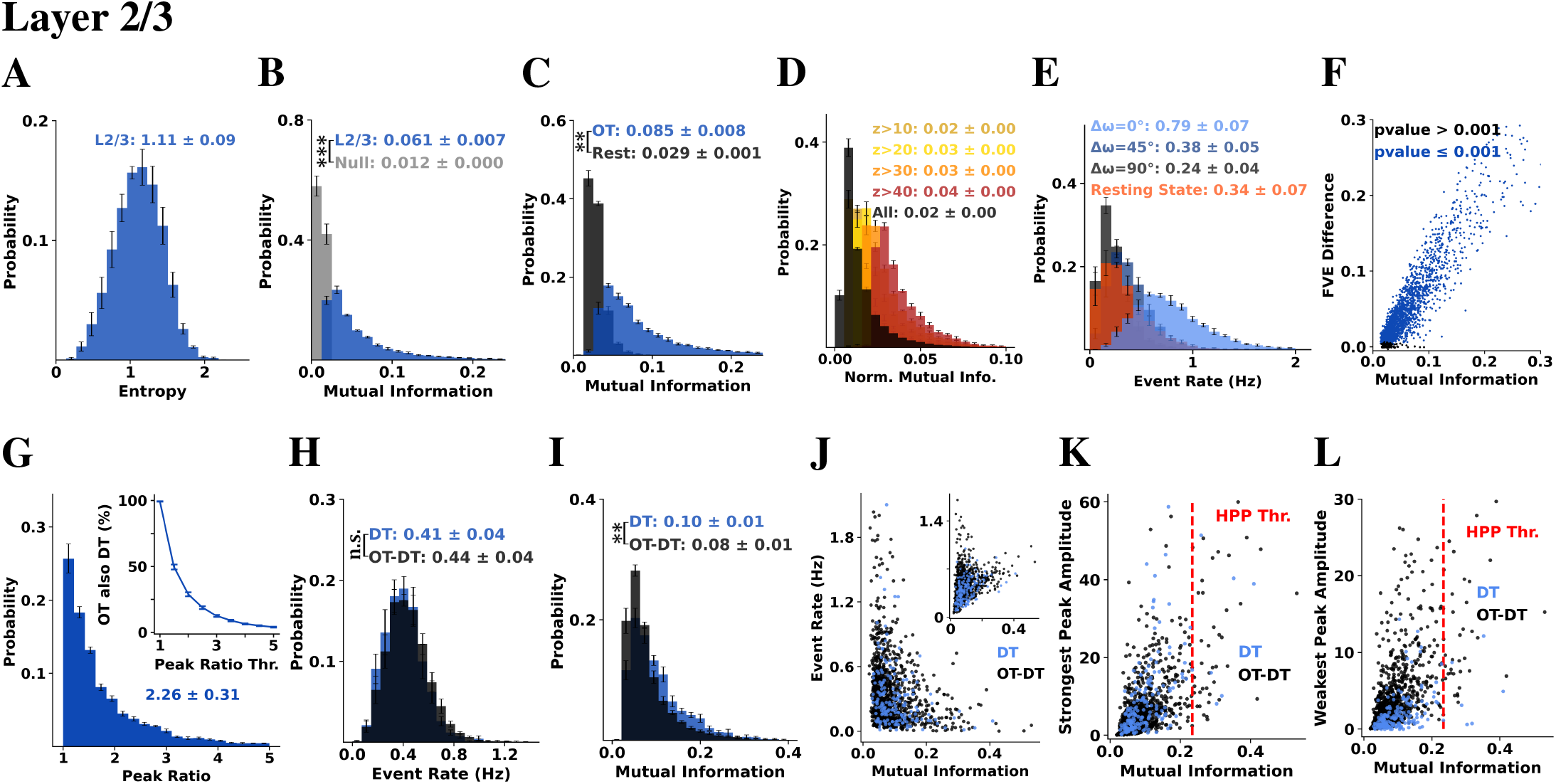
An overview of the mutual information between the event rate per segment (ERPS) of L2/3 neurons and the angle of the stimulus being presented, across all visually responsive neurons, considering different subpopulations (e.g., OT, DT, OT-DT, Rest). HPP neurons are those with the highest mutual information between their ERPS during visual stimulation and the angle of the stimulus (included here but not explicitly shown). **A:** Histogram of the entropy of the ERPS of L2/3 neurons. **B:** Histogram of the mutual information between the ERPS and the stimulus for all L2/3 neurons (blue) contrasted with that of the null (gray), which was computed by randomly shuffling the ERPS time series of each neuron 1,000 times and computing the mean mutual information across iterations. **C:** Histograms of the mutual information of the ERPS and the stimulus for orientation-tuned (OT; blue) and rest (Rest; black) L2/3 neurons. **D:** Histograms of the distributions of normalized mutual information between the ERPS and the stimulus of L2/3 neurons using different z-score thresholds (on their null MI distributions). **E:** Histogram of the ERPS for orientation-tuned L2/3 neurons during segments of a specific offset Δ*ω*. The offset Δ*ω* denotes the difference between the orientation of the stimulus being presented during a segment and the orientation preference *ϕ*_0_ of the neuron. To compute Δ*ω*, the orientation preferences of the neurons were binned with a bin size of 22.5°. The event rate under resting state conditions is included for reference. We observe significantly higher event rates when neural preference matches the stimulus direction (i.e., Δ*ω* = 0) and lower event rates when the presented stimulus is orthogonal to the cells’ preference. **F:** The scatterplot reports for each neuron of an example mouse: its mutual information with the stimulus (x-axis), its FVE difference of the von Mises fit (y-axis), color-coding the pvalue of its von Mises fit. **G:** Histogram of the peak ratio of orientation-tuned L2/3 neurons. The peak ratio represents the ratio of the amplitudes of the higher and lower peaks, derived from the two-peak von Mises fit. A higher peak ratio indicates increased direction selectivity. Inset: Percentage of direction-selective orientation-tuned neurons as a function of the peak ratio threshold. In the following figures, a neuron is defined as direction-tuned (DT), if its peak ratio is above the threshold of 3. The orientation-tuned cells that do not meet this criterion are labeled as OT-DT. **H:** Histogram of the ERPS (Hz) of L2/3 DT (blue) and OT-DT (black) neurons as computed during the stimulus presentation period. When expressed as a percentage of spiking frames, the mean event rate is ~6%. **I:** Histogram of the mutual information with the stimulus angle of DT neurons (blue) and OT-DT (black) ones. DT neurons exhibit slightly higher mutual information than OT-DT. Figs. **J-L** correspond to an example mouse. **J:** Scatterplot of the mutual information (x-axis) and the event rate during the *resting-state period* (y-axis) of L2/3 orientation-tuned neurons. Blue represents direction-tuned (DT) neurons, while black represents orientation-tuned (OT) neurons that are not DT. Inset: Same, but considering the event rate during the entire stimulus presentation period. **K:** Scatterplot of the mutual information (x-axis) and the amplitude of the stronger of the two peaks in the von-Mises fit (y-axis). The dashed red line indicates the threshold on the MI value for HPP neurons. **L:** Same as K, but for the weaker peak. HPP neurons (i.e., the top 50 neurons with the highest mutual information of ERPS and the stimulus) exhibit a broad distribution of orientation and direction tuning parameters; while some are strongly tuned to orientation, others show weaker tuning, yet still convey information about the stimulus. The mean ± standard deviation of the sample means across mice (n=5) are reported in the histograms’ insets, while error bars correspond to the standard error of the mean (SEM) across mice (n=5). P-values: “**”: < 0.01; “***”: < 0.001, and “n.s.”: non statistically significant. The highest p-value obtained from the permutation of means, the Welch’s t-test, and the ANOVA F-test is considered for the level-of-significance.

**Fig. 4:**
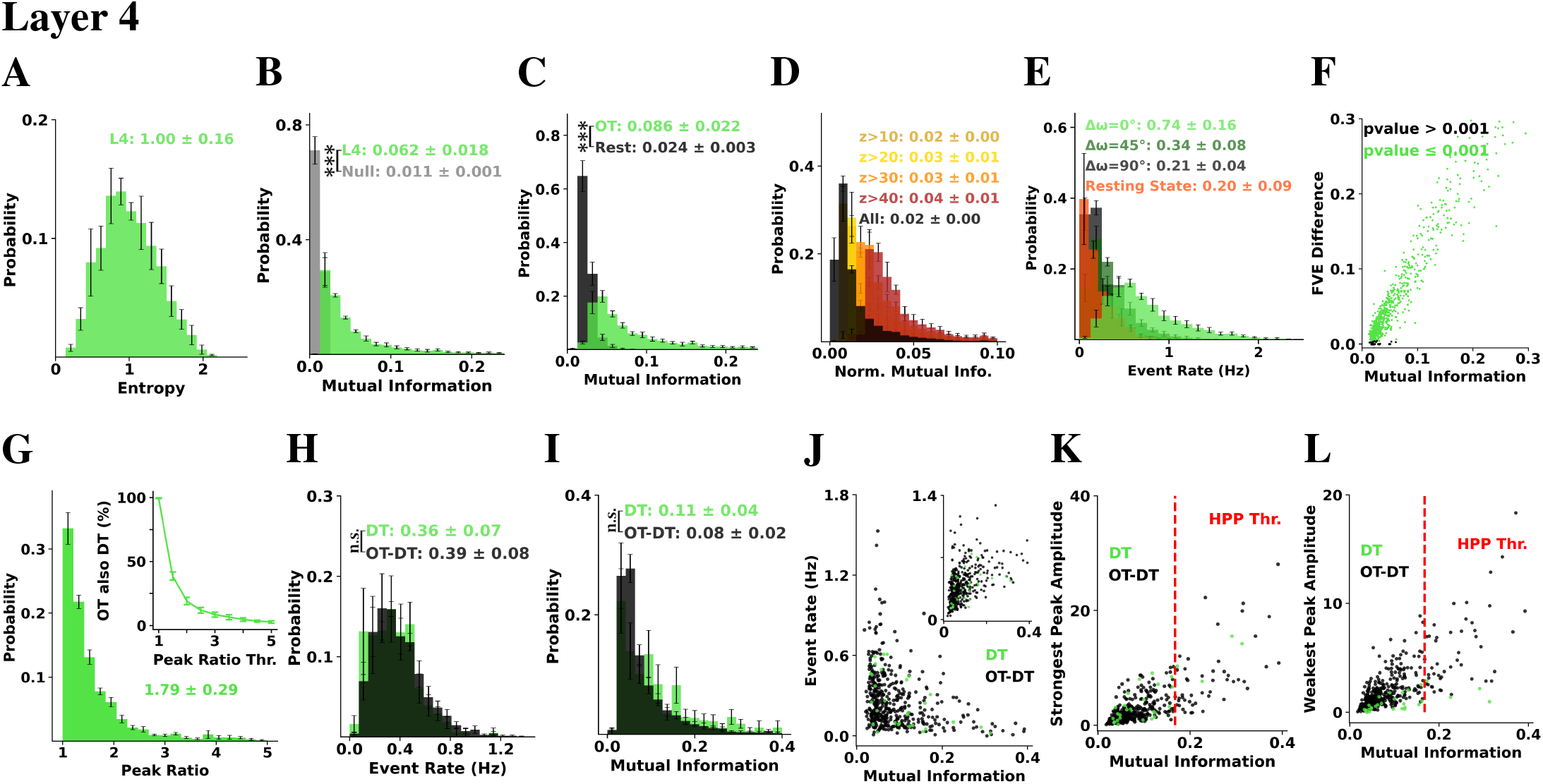
An overview of the mutual information between the event rate per segment (ERPS) of L4 neurons and the angle of the stimulus being presented, across all visually responsive neurons, considering different subpopulations (e.g., OT, DT, OT-DT, Rest). HPP neurons are those with the highest mutual information between their ERPS during visual stimulation and the angle of the stimulus (included here but not explicitly shown). **A:** Histogram of the entropy of the ERPS of L4 neurons. **B:** Histogram of the mutual information between the ERPS and the stimulus for all L4 neurons (green) contrasted with that of the null (gray), which was computed by randomly shuffling the ERPS time series of each neuron 1,000 times and computing the mean mutual information across iterations. **C:** Histograms of the mutual information of the ERPS and the stimulus for orientation-tuned (OT; green) and rest (Rest; black) L4 neurons. **D:** Histograms of the distributions of normalized mutual information between the ERPS and the stimulus of L4 neurons using different z-score thresholds (on their null MI distributions). **E:** Histogram of the ERPS for orientation-tuned L4 neurons during segments of a specific offset Δ*ω*. The offset Δ*ω* denotes the difference between the orientation of the stimulus being presented during a segment and the orientation preference *ϕ*_0_ of the neuron. To compute Δ*ω*, the orientation preferences of the neurons were binned with a bin size of 22.5°. The event rate under resting state conditions is included for reference. We observe significantly higher event rates when neural preference matches the stimulus direction (i.e., Δ*ω* = 0) and lower event rates when the presented stimulus is orthogonal to the cells’ preference. **F:** The scatterplot reports for each neuron of an example mouse: its mutual information with the stimulus (x-axis), its FVE difference of the von Mises fit (y-axis), color-coding the pvalue of its von Mises fit. **G:** Histogram of the peak ratio of orientation-tuned L4 neurons. The peak ratio represents the ratio of the amplitudes of the higher and lower peaks, derived from the two-peak von Mises fit. A higher peak ratio indicates increased direction selectivity. Inset: Percentage of direction-selective orientation-tuned neurons as a function of the peak ratio threshold. In the following figures, a neuron is defined as direction-tuned (DT), if its peak ratio is above the threshold of 3. The orientation-tuned cells that do not meet this criterion are labeled as OT-DT. **H:** Histogram of the ERPS (Hz) of L4 DT (green) and OT-DT (black) neurons as computed during the stimulus presentation period. When expressed as a percentage of spiking frames, the mean event rate is ~6%. **I:** Histogram of the mutual information with the stimulus angle of DT neurons (green) and OT-DT (black) ones. DT neurons exhibit slightly higher mutual information than OT-DT. Figs. **J-L** correspond to an example mouse. **J:** Scatterplot of the mutual information (x-axis) and the event rate during the *resting-state period* (y-axis) of L4 orientation-tuned neurons. Green represents direction-tuned (DT) neurons, while black represents orientation-tuned (OT) neurons that are not DT. Inset: Same, but considering the event rate during the entire stimulus presentation period. **K:** Scatterplot of the mutual information (x-axis) and the amplitude of the stronger of the two peaks in the von-Mises fit (y-axis). The dashed red line indicates the threshold on the MI value for HPP neurons. **L:** Same as K, but for the weaker peak. HPP neurons (i.e., the top 50 neurons with the highest mutual information of ERPS and the stimulus) exhibit a broad distribution of orientation and direction tuning parameters; while some are strongly tuned to orientation, others show weaker tuning, yet still convey information about the stimulus. The mean ± standard deviation of the sample means across mice (n=5) are reported in the histograms’ insets, while error bars correspond to the standard error of the mean (SEM) across mice (n=5). P-values: “**”: < 0.01; “***”: < 0.001, and “n.s.”: non statistically significant. The highest p-value obtained from the permutation of means, the Welch’s t-test, and the ANOVA F-test is considered for the level-of-significance.

The **visually responsive neurons** are defined as those that carry information about the stimulus. To do so, we used statistical tests based on mutual information between neuronal firing and stimulus. Note that this potentially excludes neurons that significantly respond to the stimulus directions homogeneously versus baseline. To adopt a more conservative criterion for visual responsiveness, we additionally include a control for each neuron based on the firing events per segment estimated *during the resting state* (i.e., in the absence of stimulus presentation). For each neuron, we then compute a second control mutual information (MI) value between the event rate per segment (ERPS) time series during the resting state and the sequence of stimulus angles, following the same methodology and number of repetitions as for the control MI based on randomly shifted ERPS during stimulus presentation. We define a neuron to be **visually responsive**, if its MI is above the 99.9% of both its corresponding null distribution of MI values. The percentage of visually responsive neurons appears consistent with that reported in [7] (although their results were obtained using different statistical tests). All plots in this paper focus *exclusively* on visually responsive neurons.

### Orientation- and Direction-Tuned Neurons

The neurons’ orientation and direction tuning were estimated as in [37]. Briefly, responses to a dynamic stimulus of pink noise with coherent orientation and motion (i.e., Monet video) were fit with a *two-peak von Mises* function. Cells that satisfy a dual threshold for the *fraction of variance explained* (*FVE*), namely, the difference between the original fit FVE and the median

FVE across all 1000 shuffled fits *>*2.5% and significance calculated by permutation *p* ≤ 0.001 (see Appendix for more details) are defined ***orientation-tuned (OT)***. All FVE values (one per neuron) that we report on the graphs of this work correspond to this *difference* in fraction variable explained between the *original* fit (FVE) and the median FVE in all 1000 *shuffled* fits. In our sample, for an example mouse, ~58% of L4-neurons (469 out of 805 visually responsive neurons) and ~58% (1,188 out of 2,032 visually responsive neurons) of L2/3-neurons were orientation-tuned (OT) to the stimulus using these criteria, consistent with [37]. All orientation-tuned units were then sorted “cyclically” into 128 polar angle bins according to the preferred direction of the *larger* amplitude von Mises peak.

Note that statistically significant orientation-tuned (OT) neurons (as defined here) contain some information about directionality when the two peaks are not equal. A higher peak indicates an increase in direction selectivity. For each orientation-tuned neuron, the peak ratio was defined as the *ratio* between the *amplitudes of the higher* and *lower peaks* in its tuning curve. An OT neuron is defined as ***direction-tuned (DT)***, if its peak ratio is *above the threshold of 3. Only* orientation-tuned neurons, not direction-tuned ones, are indicated by the **“OT-DT”** (i.e., OT *minus* DT).

L2/3 neurons demonstrate greater direction selectivity compared to L4 neurons (Table I). Across both layers, DT neurons are significantly fewer than non direction-tuned OT neurons, for any direction selectivity threshold set at 2 or higher, as illustrated in Figs. 3G and 4G.

**Table 1.** Percentages and absolute counts (in parentheses) of DT (direction-tuned), OT-DT (orientation-tuned, but not direction-tuned) and Rest neurons within layers L4 and L2/3 in V1. Only visually responsive V1 neurons are considered. We report the mean and standard deviation across five mice (n=5).

|  | Layer 4 | Layer 2/3 |
| --- | --- | --- |
| **DT** | 5% ± 1% (31 ± 7) | 7% ± 1% (185 ± 15) |
| **OT-DT** | 53% ± 11% (391 ± 179) | 50% ± 4% (1,290 ± 399) |
| **Rest** | 42% ± 11% (272 ± 92) | 43% ± 4% (1,086 ± 242) |

As expected, statistically significant orientation-tuned neurons exhibit significantly higher MI than the rest (Figs. 3C & 4C). A substantial percentage of neurons has statistically significant normalized mutual information (Figs. 3D & 4D). An exclusively direction-selective unit that *always* fires in response to a specific direction and remains silent otherwise exhibits lower mutual information with the stimulus (~0.34) compared to a “perfectly” orientation-selective unit (~ 0.54). However, when the firing of this theoretical orientation-selective unit is adjusted to match that of the direction-selective unit (1/16), by reducing its firing probability for the preferred orientation to 1/2, the mutual information of this unit decreases to ~0.304, falling below that of the direction-selective neuron. This is consistent with the experimental results, which show that both the DT and OT-DT populations exhibit an event rate of approximately 6%, when expressed as a percentage of spiking frames (Figs. 3H & 4H), yet DT neurons demonstrate slightly higher mutual information (Figs. 3I & 4I), suggesting that direction-selective neurons may encode more information about the stimulus at low firing event rates, facilitating more efficient neural processing of the stimulus.

### HPP neurons

We examined the L2/3 and L4 neurons with respect to their tuning characteristics, event rate, and MI (Fig. 3 & 4) and focused on neurons with high MI. This distinction represents a continuum rather than a strict dichotomy (Figs. 3J, 3K, 3L,4J, 4K, 4L). Neurons with high predictive power (HPP) with stimulus are defined here as the neurons *ranked highest in the layer*, in terms of their nMI with the angle of the presented stimulus. To highlight the main trend, we selected the *top 50 neurons* in terms of their MI (Fig. 3 & 4) and compared them with the **orientation-tuned neurons excluding the HPP (OT-HPP)** vs. the nonorientation-tuned ones (Rest). All identified HPP neurons are orientation-tuned. HPP neurons exhibit a broad distribution of orientation and direction tuning parameters; while some are strongly tuned to orientation, others show weaker tuning, yet still convey information about the stimulus (Figs. 3K, 3L, 4K, 4L).

If we relax the threshold to include the top 10% of neurons (corresponding to the start of the plateau in decoding performance, Fig 5C), the trends observed below persist.

**Fig. 5:**
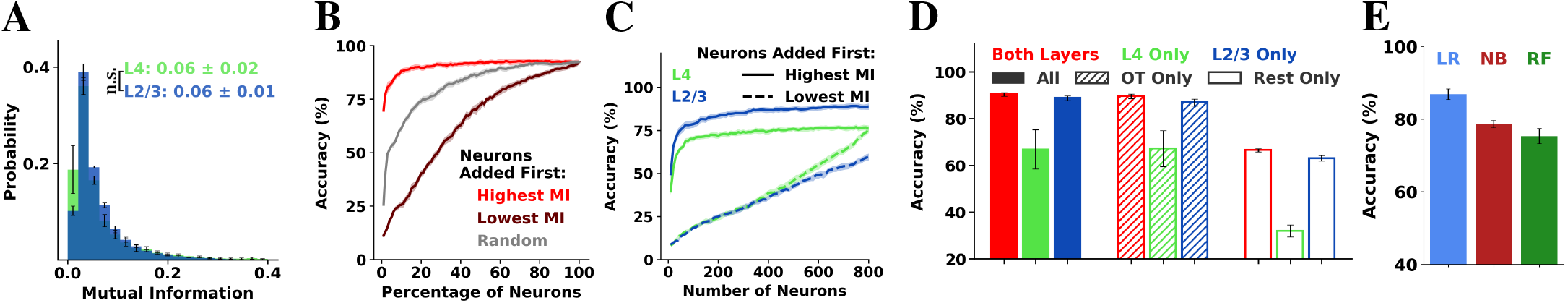
Multi-class classification for predicting the angle of the stimulus. **A**: Histograms of the mutual information of L4 (L2/3) neurons, presented in green (blue), respectively. The difference is non-significant (n.s.). **B:** Accuracy of Logistic Regression for predicting the angle of the stimulus as a function of the percentage of neurons employed (features) for an example mouse. Specifically, all V1 neurons were sorted by the normalized mutual information of their event rate per segment with the stimulus. The x-axis indicates the X% of neurons selected for the prediction of the stimulus (in descending order of MI for the red line, in ascending order of MI for the brown line). The input of the classifier were the event rates per segment of each selected neuron, standardized as to have a mean of 0 and a standard deviation of 1. We performed 5-fold cross validation. We report the mean (solid lines) and SEM (shaded region) of the accuracy across folds for an example mouse. For the null (gray line), neurons are selected in a random order. Note that the accuracy for highest MI neurons added first (red line) plateaus for all mice after 20%, indicating that no significant information regarding the stimulus can be gained by adding more neurons. **C:** Same as B, but using as features only L4 (L2/3) neurons, depicted in green (blue), respectively. Here, the x-axis indicates the number of neurons selected. We report the accuracy in the case that we select neurons in descending (solid line) and ascending (dashed line) order of MI. **D:** Accuracy of Logistic Regression in predicting the angle of the stimulus being presented during a segment using as features the ERPS of different neuronal subpopulations: all neurons (solid), only OT neurons (dashed), and only Rest neurons (white). Red indicates models using neurons from both L4 and L2/3; green and blue indicate models using only L4 or only L2/3 neurons, respectively. We performed 5-fold cross validation and computed the mean accuracy across folds per mouse. The bars correspond to the mean and the error bars to the SEM of the accuracy across mice (n=5). **E:** Accuracy of Logistic Regression (LR), Naïve Bayes (NB), and Random Forest (RF) models in predicting the angle of the stimulus using as features the ERPS of the OT L2/3 neurons.

**Fig. 6:**
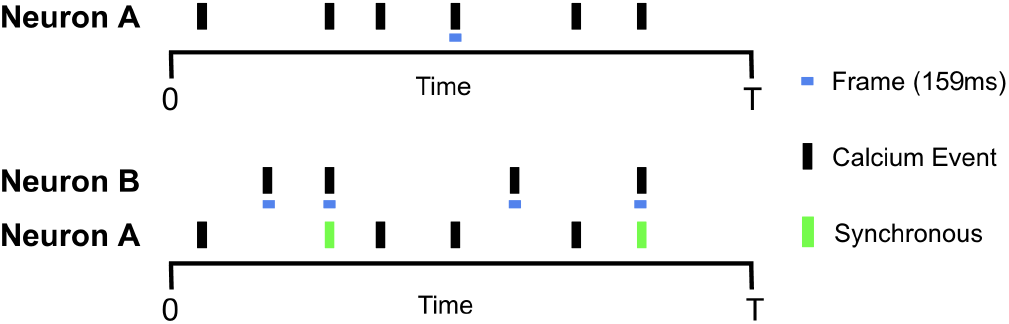
Spike Time Tiling Coefficient (STTC) Diagram of the STTC estimation, adapted from [32]. The temporal relationship between calcium events of neurons A and B is analyzed within a time window of ±ΔT (here, ΔT = 0), capturing coincident calcium events. Vertical lines denote the calcium events of neurons. The green bars mark the calcium events of A that fall within Δt (“tile”, blue bar) of calcium events of B. See section V for more details.

We found that HPP neurons exhibit higher event rates than OT-HPP neurons, which, in turn, have higher event rates than rest, as estimated throughout the period under stimulus presentation (Figs. 7A & 8A). The distribution of the event rates per segment during different periods is demonstrated in Figs. 3E and 4E. For segments where the stimulus direction is approximately at 45 degrees away from the cell’s preferred direction, the mean event rate is 0.38 Hz and 0.34 Hz compared to 0.79 Hz and 0.74 Hz for segments where the stimulus direction matches the cell’s preference in L2/3 and L4, respectively (Figs. 3E & 4E). Moreover, the event rate per segment in response to orthogonal stimuli can be low, sometimes even lower than during the resting state, indicating inhibition of firing in the presence of stimuli for neurons with orthogonal preferences for the presented stimulus, consistent with previous studies [38].

**Fig. 7:**
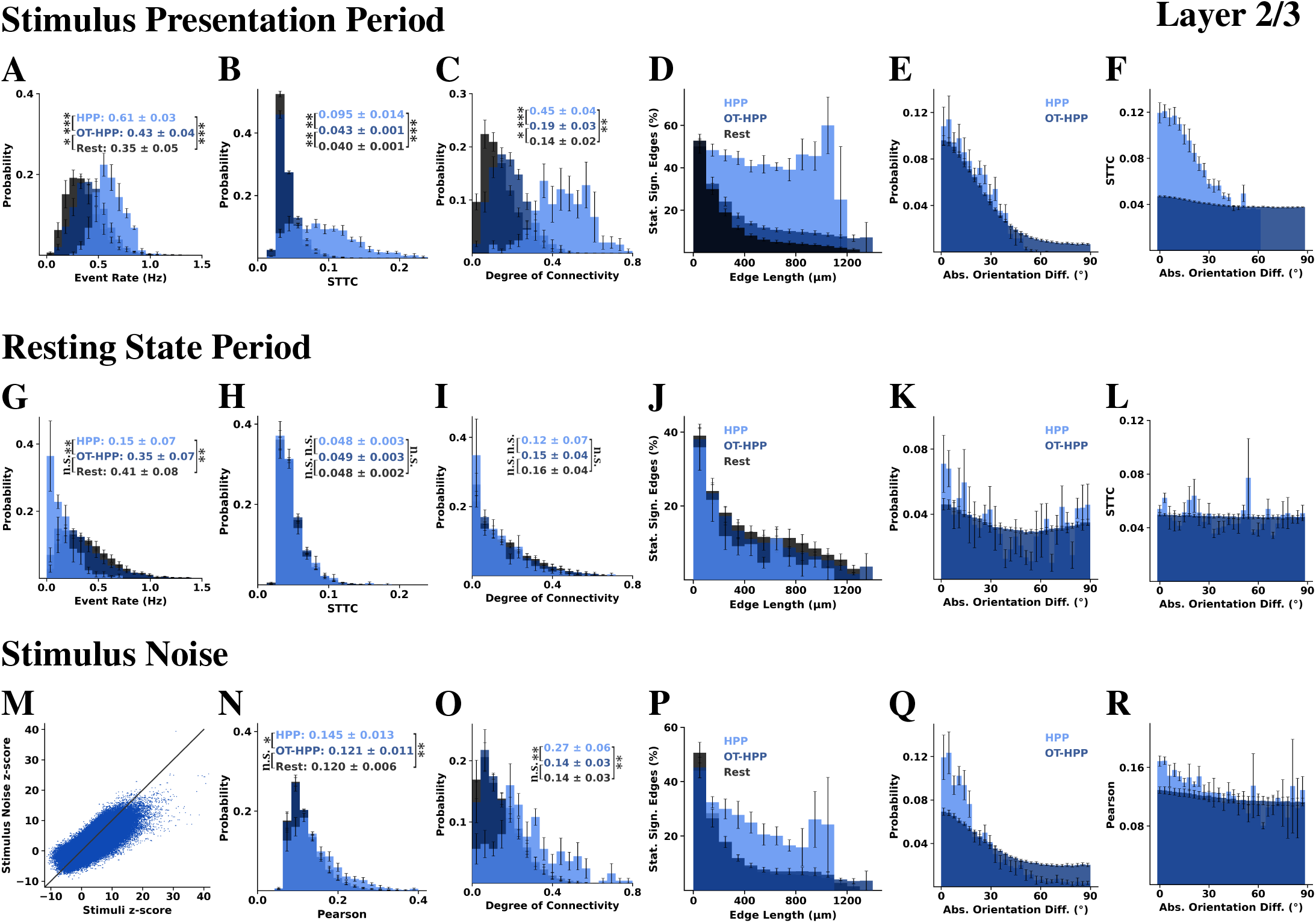
Functional Connectivity Analysis. L2/3-HPP neurons have distinct functional structure *under stimulus presentation* (top and bottom rows) compared to the structure *under resting state* (middle row). The first and third rows demonstrate the functional connectivity driven by stimulus (top) vs. the functional connectivity based on the noise correlations (bottom). Even removing the main stimulus effect (tuning) the stimulus correlation structure remains. **A & G:** Histograms of the event rates during the entire stimulus presentation and resting state periods, respectively. HPP neurons exhibit higher event rates than the other two sub-populations under stimulus presentation, however, the trend reverses during resting state conditions. **B, H & N:** Histograms of the correlation weight of statistically significant (z-score*>*4) positive functional correlations in L2/3, among pairs of neurons that belong in the same category, computed using STTC on the eventograms obtained during the stimulus presentation period (Fig. B), using STTC on the eventograms obtained at resting state (Fig. H), and using Pearson correlation on the stimulus noise (Fig. N),. **C, I & O:** Histograms of the degree of functional connectivity. Considering here exclusively the intra-group functional connectivity, i.e., between neurons of the same type, namely HPP, OT-HPP, and rest, we found that HPP neurons exhibit a higher intra-group degree of connectivity compared to the other two populations during stimulus presentation. Interestingly, when we consider exactly these three neuronal subpopulations and examine their functional connectivity within each subpopulation *under resting state conditions, there are no statistically significant differences*. **D, J & P:** Percentage of statistically significant edges (z-score*>*4) as a function of distance between neuronal pairs, plotted in bins of 100*µm*. The percentage is computed by taking as a denominator the number of all possible edges that could form at that distance between neurons of the same category (namely, HPP, OT-HPP and Rest). Results from all edges within each animal are averaged, marking the high predictive power (HPP) neurons in light blue, the OT-HPP in dark blue, and the remaining in black. Bins with a single observed value were excluded. Error bars correspond to SEM across mice. **E, K & Q:** Histogram of the difference in orientation preference among HPP (light blue) and OT-HPP (dark blue) neuronal pairs with statistically significant functional correlations (z-score *>* 4). Orientation difference between neuronal pairs was estimated as the minimum absolute difference of their strongest amplitude angles computed in the circular [0,180) space (zero identified to 180 degrees); the orientation-difference range is, therefore [0, 90] degrees. Error bars indicate the SEM across mice. **F, L & R:** Weight of the pairwise correlation of edges among HPP (light blue) and OT-HPP (dark blue) neuronal pairs as a function of their absolute orientation difference. The correlations were computed with STTC during stimulus presentation (Fig. F) and at resting state (Fig. L), while Pearson correlation was employed for the stimulus noise (Fig. R). Error bars indicate the SEM across mice. **M**: Scatterplot of the z-score of the STTC as computed on the eventogram of the stimulus presentation period (x-axis) and the z-score of the Pearson correlation computed on the noise signal of the stimulus presentation period (y-axis) of all intra-L2/3 neuronal pairs for an example mouse. P-values: “*”: < 0.05; “**”: < 0.01; “***”: < 0.001, and “n.s.”: non statistically significant. The highest p-value obtained from the permutation of means, the Welch’s t-test, and the ANOVA F-test is considered for the level-of-significance.

**Fig. 8:**
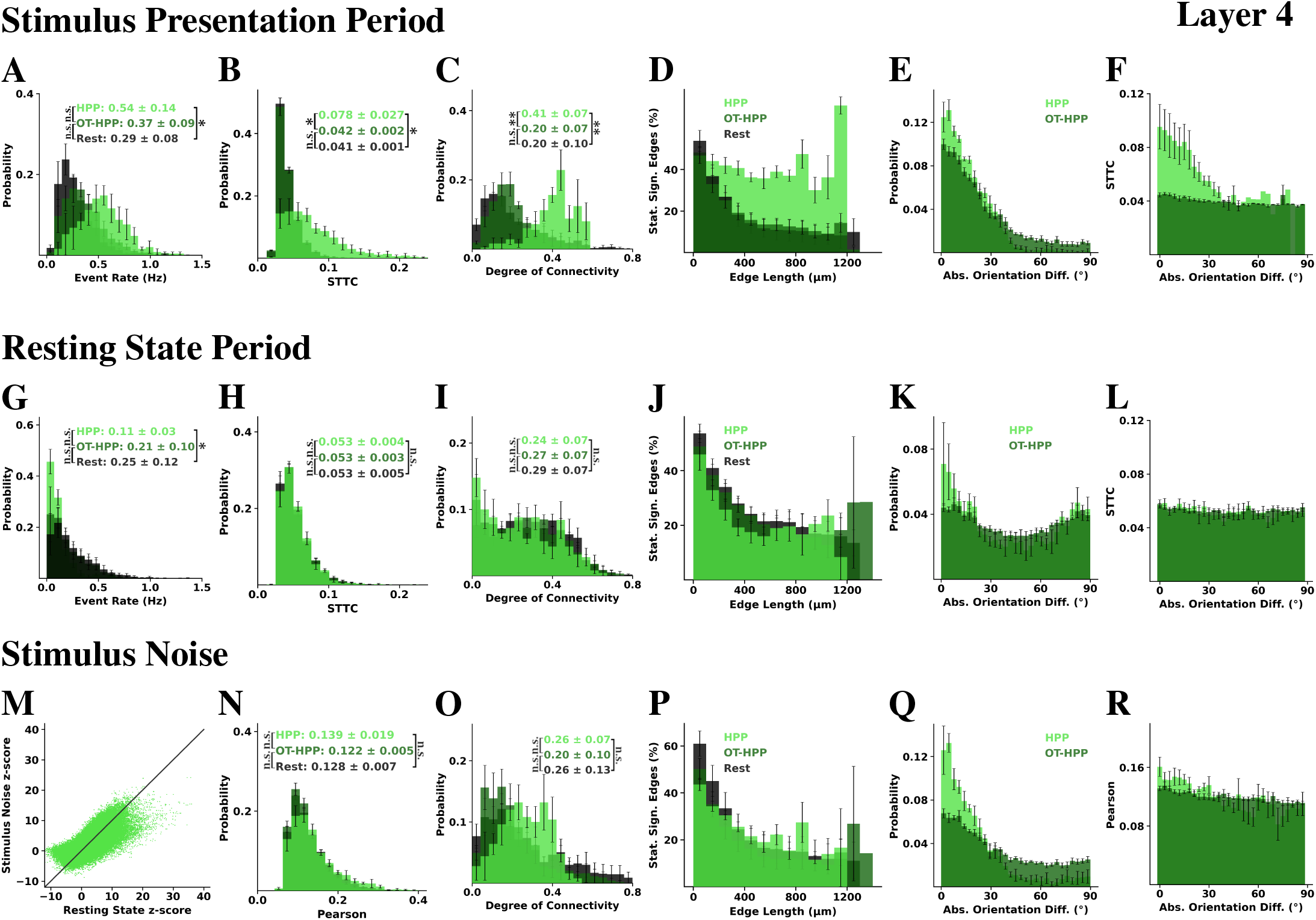
Functional Connectivity Analysis. L4-HPP neurons have distinct functional structure *under stimulus presentation* (top and bottom rows) compared to the structure *under resting state* (middle row). The first and third rows demonstrate the functional connectivity driven by stimulus (top) vs. the functional connectivity based on the noise correlations (bottom). Even removing the main stimulus effect (tuning) the stimulus correlation structure remains. **A & G:** Histograms of the event rates during the entire stimulus presentation and resting state periods, respectively. HPP neurons exhibit higher event rates than the other two sub-populations under stimulus presentation, however, the trend reverses during resting state conditions. **B, H & N:** Histograms of the correlation weight of statistically significant (z-score*>*4) positive functional correlations in L4, among pairs of neurons that belong in the same category, computed using STTC on the eventograms obtained during the stimulus presentation period (Fig. B), using STTC on the eventograms obtained at resting state (Fig. H), and using Pearson correlation on the stimulus noise (Fig. N),. **C, I & O:** Histograms of the degree of functional connectivity. Considering here exclusively the intra-group functional connectivity, i.e., between neurons of the same type, namely HPP, OT-HPP, and rest, we found that HPP neurons exhibit a higher intra-group degree of connectivity compared to the other two populations during stimulus presentation. Interestingly, when we consider exactly these three neuronal subpopulations and examine their functional connectivity within each subpopulation *under resting state conditions, there are no statistically significant differences*. **D, J & P:** Percentage of statistically significant edges (z-score*>*4) as a function of distance between neuronal pairs, plotted in bins of 100*µm*. The percentage is computed by taking as a denominator the number of all possible edges that could form at that distance between neurons of the same category (namely, HPP, OT-HPP and Rest). Results from all edges within each animal are averaged, marking the high predictive power (HPP) neurons in light blue, the OT-HPP in dark blue, and the remaining in black. Bins with a single observed value were excluded. Error bars correspond to SEM across mice. **E, K & Q:** Histogram of the difference in orientation preference among HPP (light blue) and OT-HPP (dark blue) neuronal pairs with statistically significant functional correlations (z-score *>* 4). Orientation difference between neuronal pairs was estimated as the minimum absolute difference of their strongest amplitude angles computed in the circular [0,180) space (zero identified to 180 degrees); the orientation-difference range is, therefore [0, 90] degrees. Error bars indicate the SEM across mice. **F, L & R:** Weight of the pairwise correlation of edges among HPP (light blue) and OT-HPP (dark blue) neuronal pairs as a function of their absolute orientation difference. The correlations were computed with STTC during stimulus presentation (Fig. F) and at resting state (Fig. L), while Pearson correlation was employed for the stimulus noise (Fig. R). Error bars indicate the SEM across mice. **M**: Scatterplot of the z-score of the STTC as computed on the eventogram of the stimulus presentation period (x-axis) and the z-score of the Pearson correlation computed on the noise signal of the stimulus presentation period (y-axis) of all intra-L4 neuronal pairs for an example mouse. P-values: “*”: < 0.05; “**”: < 0.01; “***”: < 0.001, and “n.s.”: non statistically significant. The highest p-value obtained from the permutation of means, the Welch’s t-test, and the ANOVA F-test is considered for the level-of-significance.

**Fig. 9:**
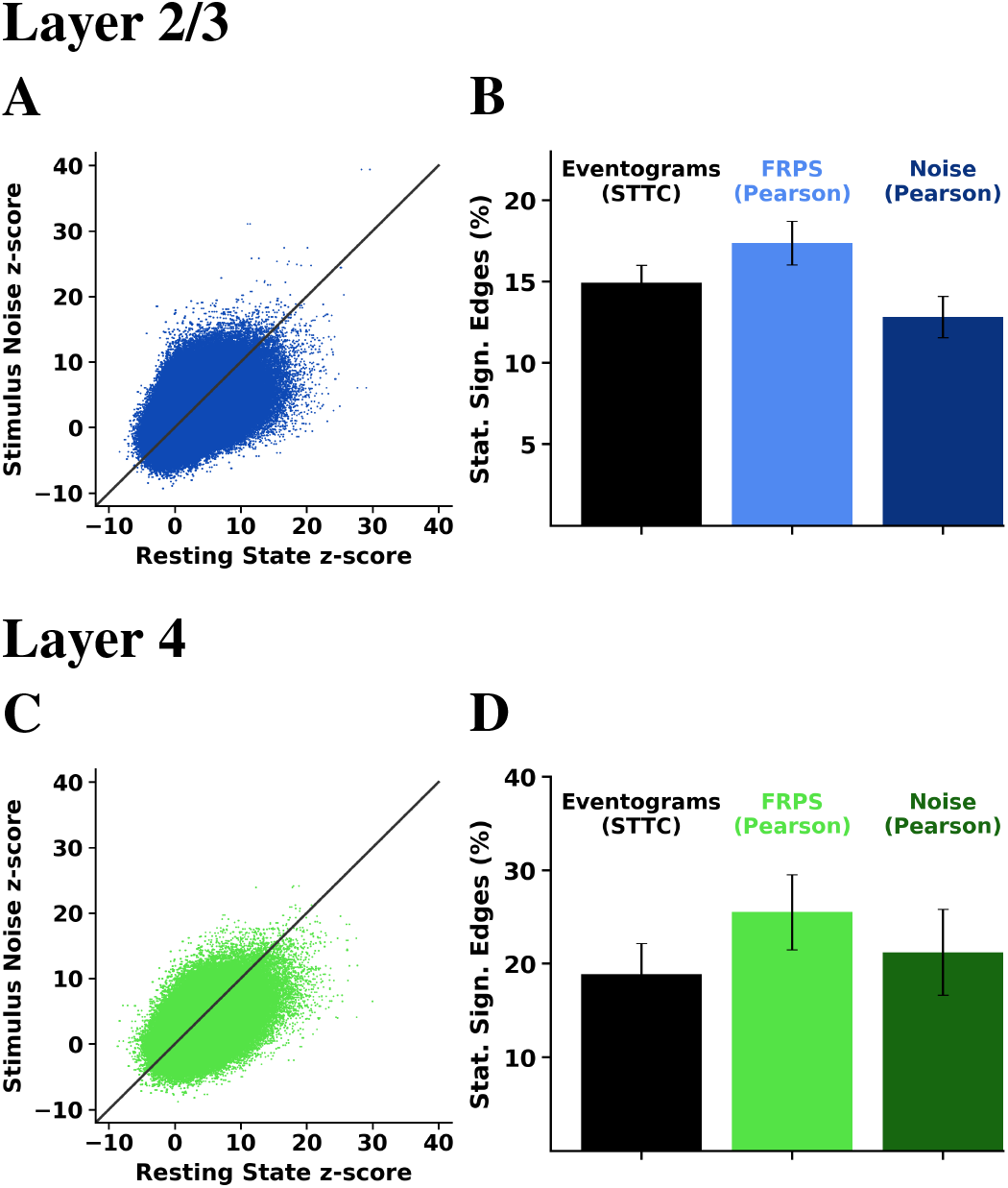
**A:** Scatterplot of the z-score of the STTC as computed on the eventogram of the resting state period (x-axis) and the z-score of the Pearson correlation computed on the noise signal of the stimulus presentation period (y-axis) of all intra-L2/3 neuronal pairs for an example mouse. **B:** Percentage of statistically significant (z-score*>*4) intra-L2/3 pairs when computed on the eventograms of the stimulus presentation period with STTC (black), on the ERPS timeseries of the stimulus presentation period using Pearson’s correlation (light blue) and on the noise signal using Pearson’s correlation (dark blue). **C & D:** Same as A & B, but considering only intra-L4 neuronal pairs.

## IV. Decoding the stimulus direction

The distributions of the MI of the ERPS of L2/3 and L4 neurons with the stimulus angle do not exhibit significant differences (Fig. 5A). To examine the role of L4 and L2/3 V1 neurons in decoding the stimulus angle, we also used various models for multi-class classification. Specifically, we employed Logistic Regression under 5-fold cross-validation and estimated the mean accuracy using as features the event rate per segment of neurons (Fig. 5). We found that the 20% highest ranking neurons in terms of MI with the stimulus are enough to reach an accuracy of around 90% (Fig. 5B, red line). This is observed in both L4 and L2/3 (Fig. 5C). Furthermore, using L2/3 neurons as input yields better predictions than using the same number of L4 neurons. Interestingly, when selecting the lowest-ranking 80% of neurons, the performance does not reach the same level (Fig. 5B, brown line). As expected, when the entire neuronal population is utilized, the decoding performance (on average) reaches its highest value (consistent with previous findings [5], [6]). However, a nearly identical accuracy is achieved using only the OT neurons (Fig. 5D, average accuracy across mice) highlighting the redundancy in firing activity. We employed the Naive Bayes classifier for the same task–a decoder that assumes *independence* of the neurons given the stimulus angle. We found that its performance had considerably worse accuracy–approximately 10% lower than Logistic Regression (Fig. 5E). This finding is consistent with previous work [5] and suggests the potential impact of the inter-neuronal dependencies on decoding the stimulus direction. All classifiers were implemented using scikit-learn, and default parameters were utilized. No additional hyperparameter tuning was performed. The dataset was preprocessed using standard normalization techniques.

## V. Functional Connectivity

To identify functional connectivity patterns within and across the recorded layers, we applied the Spike Time Tiling Coefficient (STTC), a pairwise functional connectivity measure that exhibits several advantages over several other correlation measures (e.g., depends less on the firing rate than Pearson correlation) [32].

### Estimation of pairwise temporal correlation

To quantify the temporal correlation between the firing events of a neuronal pair A and B, we employed their calcium eventograms and estimated the **STTC** weight as follows:

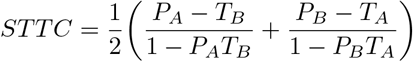

where *T*_*A*_ is the proportion of the recording duration that corresponds to firing events of neuron A, *P*_*A*_ the proportion of firing events of neuron A *synchronous* (i.e., within the same 158.7 ms frame) with firing events of neuron B, and likewise for *T*_*B*_ and *P*_*B*_. The STTC weight (correlation) takes values in [-1, 1]. To determine whether there is a *statistically significant functional connection between two neurons*, their STTC value was compared to a null distribution of STTC values calculated by circularly shifting their calcium eventogram time series by a random number, 500 times, yielding a z-score that determines the *level of significance*. We consider that two neurons are *functionally connected* when their z-score is *above 4*.^4^ Note that given the temporal kinetics of calcium imaging and the frame duration in our datasets (*≈* 158.7 ms), the duration of a single frame is large compared to the neuron communication time (few ms).

Considering exclusively the functional *intra-layer* connectivity *between neurons of the same type* (e.g., only between HPP neurons or between Rest neurons), in the *same layer*, we found that, under visual stimulation, HPP neurons have stronger statistically significant correlations (see Figs. 7B & 8B) and exhibit a higher degree of connectivity within their HPP group than members of the other two populations within their own groups (see Figs. 7C & 8C). This is not surprising. Interestingly, even removing the main stimulus effect (tuning), by subtracting the mean firing, the stimulus correlation structure is not eliminated completely, i.e., the functional connectivity driven by stimulus is consistent with the functional connectivity based on noise correlations (Figs. 7 & 8). Specifically, functional connectivity based on noise correlations was identified as follows: The stimulus noise time series was formed per neuron by subtracting from the event rate per segment time series (at segment *t*) the average event rate computed using all segments during which the stimulus was presented is the same as the one at segment *t*. That is, if 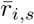 is the mean event rate of neuron *i* during presentation of stimulus *s*, the stimulus noise time series is 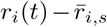, where *r*_*i*_(*t*) is the event rate of neuron *i* during segment *t* and *s* the stimulus presented at that segment (t). We then applied Pearson correlation to form the functional connectivity based on noise correlations. Thus, although stimulus-driven functional connections between HPP neurons are relatively longer than intragroup connections within OT-HPP and rest (e.g., during visual stimulation see Figs. 7D & 8D), no significant differences in connection lengths are observed among the three groups during resting state (Figs. 7J & 8J). This discrepancy between the HPP vs. control groups between conditions becomes evident also when examining the correlation strength (Figs. 7H vs. 7N, 8H vs. 8N) and the degree of intra-group connectivity within the specific subpopulations (Figs. 7I vs. 7O, 8I vs. 8O). For example, HPP neurons exhibit a higher degree of connectivity than the rest during visual stimulation, whereas all three populations show similar connectivity patterns during the resting state. This suggests that spontaneous activity patterns do not consistently reflect those induced by external stimuli.

## VI. Discussion and Future Work

We identified the visually responsive neurons and comparatively examined their overall event rate, tuning properties, and predictive power about the angle of the stimulus (Figs. 3 & 4). The distinction reflects a continuum rather than a strict dichotomy. HPP neurons display a wide range of orientation and direction tuning properties; some are sharply tuned to orientation, while others are more weakly tuned, yet still carry information about the stimulus (Figs. 3K, 3L, 4K, 4L). HPP neurons exhibit elevated calcium event rates during stimulus presentation (Fig. 7A & 8A), as expected. However, interestingly, during resting-state, this event rate pattern reverses (Figs. 7G & 8G). This suggests that they are less strongly modulated by the internal state. During stimulus presentation, the architecture of functional connectivity among HPP neurons is highly distributed and denser compared to connectivity between other control groups (Figs. 7C & 8C). In contrast, during resting-state, their connectivity does not differ significantly in structure from that of the control groups (Figs. 7H-7L, 8H-8L). Although HPP neurons work together as a group under stimulus conditions, they do not exhibit strong synchronization as measured by functional connectivity under resting-state conditions. This suggests that spontaneous patterns of activity do not always recapitulate stimulus-induced activation patterns.

We also evaluated the decoding ability of various neuronal populations using Logistic Regression, Naive Bayes, and Random Forests. Relatively *small sub-groups* of HPP neurons in the population *carry most of the predictive power* and alone can reach the prediction accuracy demonstrated by the whole population (Fig. 5B). Moreover, the observation that the top L2/3 neurons, ranked by MI, exhibit higher predictive power than an equal number of top-ranked L4 neurons, suggests that V1 processing enhances stimulus detection (e.g., Figs. 5C, 5D), supporting the view of L2/3 as the “output” of V1, in contrast to L4, which primarily functions as its “input” layer. Our analysis is subject to certain limitations, including the relatively low temporal resolution of the calcium signal and the use of the Monet stimulus in place of naturalistic stimuli. Despite these constraints, the findings reveal intriguing patterns that merit further investigation. Several key questions remain unanswered—chief among them, how dynamically interacting neuronal populations coordinate to encode distinct features of a stimulus, and what the precise relationship is between spontaneous and stimulus-driven activity patterns. We propose that the activity patterns emerging from architecturally connected neuronal ensembles constitute a kind of “vocabulary space”. Deciphering the functional architecture of the cortex may help us understand how this “vocabulary space” is organized and deployed across different stimulus conditions, ultimately offering insights into the fundamental principles of cortical information processing.

## Acknowledgments

We would like to thank the members of Tolias Lab at the Department of Neuroscience at Baylor College of Medicine in Houston, Texas for performing the two-photon (mesoscope) experiments, sharing the datasets with us, and providing valuable feedback about the data. We especially acknowledge Paul G. Fahey for the discussions about the data. We are also grateful to the data scientists at the Institute of Computer Science, FORTH, namely, Nikolaos Tzanakis, Manos Koniotakis, and Vassilis Sideridis, who contributed to the implementation of several aspects of the preliminary analysis.

This work has received funding from the European Union’s Horizon 2020 research and innovation program under the Marie Skłodowska-Curie grant agreement No 101007926 as well as from the Hellenic Foundation Research Institute (HFRI) with the neuron-AD project number 04058 and neuronXnet project number 2285 (PI: Maria Papadopouli). It has been partially supported by project MIS 5154714 of the National Recovery and Resilience Plan Greece 2.0 funded by the European Union under the NextGenerationEU Program. Finally, this research was also supported by R01 NS113890, and R21 NS127299 (PI: Stelios Smirnakis).

## [Appendix

## Mouse Lines and Surgery

Five adult mice (10-12 weeks of age), expressing GCaMP6s in excitatory neurons via SLC17a7-Cre and Ai162 transgenic lines, were anesthetized and a 5mm craniotomy was placed over visual cortex as described [37]. Each mouse recovered for~ 2 weeks prior to the first experimental imaging session.

### Experimental Data Collection

The animals underwent mesoscopic two-photon imaging covering most of dorsal area V1 and nearby extrastriate cortex, while being head-fixed on a treadmill in quiet wakefulness. Images were acquired at 6.30072 Hz over a ~ 1.2×1.2 mm^2^ field of view sampling simultaneously across 4 planes corresponding to V1 layers 2 (80-210 mm), 3 (285-330 mm), 4 (400-450 mm) and 5 (500 mm). Images were preprocessed in standard fashion for motion correction and underwent automatic segmentation and deconvolution using the CNMF CaImAn algorithm [30]. The deconvolved signal was thresholded appropriately to yield calcium “eventograms” that were used for analysis. The threshold yielding calcium event rates closer to those reported in the literature [31] was selected. Neurons located less than 15mm from the periphery of the field of view (FOV) were excluded in order to avoid potential edge effects arising from incomplete correction of motion artifacts.

### Monitor Positioning and Retinotopy

Visual stimuli were presented to the left eye with a 31.1×55.3cm^2^ (h×w) monitor (resolution of 1440×2560 pixels) positioned 15cm away from the mouse eye. Pixelwise responses across a 2400×2400 *µ*m^2^ to 3000×3000 *µ*m^2^ region of interest (0.2 px/*µ*m) at 200-220*µ*m depth from the cortical surface to drifting bar stimuli were used to generate a sign map for delineating visual areas [39]. The directional trial response was measured by taking the difference in cumulative deconvolved activity at the linearly interpolated trial onset and offset time points. Trial responses per direction were modeled as a two-peak scaled von Mises function (see [37]). The two peaks share a preferred orientation, baseline, and width, but their amplitudes are fit independently. This function was fitted to minimize the mean squared error of all trial responses *across 16 directions* using the L-BFGS-B optimization algorithm [40]. Significance and goodness of fit were calculated by permutation. Trial direction labels were randomly shuffled among all trials for 1000 refits. The goodness of fit was calculated as the difference in *fraction variable explained (FVE)* between the original fit FVE and the median FVE across all 1000 shuffled fits. The p-value was calculated as the fraction of shuffled fits with a higher FVE than the original fit.

### Normalized Degree of Connectivity

For each neuron, we estimate the fraction of neurons with statistically significant functional connections (z-score *>* 4) to that neuron, per layer case, which corresponds to the normalized degree of connectivity in that layer. For example, a L4 neuron has an intra-layer degree of connectivity of 0.1 if that neuron is functionally connected with the 10% of the L4 neuronal population.

### Absolute Orientation Difference

For each pair of neurons (n1, n2), we estimate the absolute direction difference of their strongest amplitude angle w1 and w2, respectively, *ϕ* as follows:

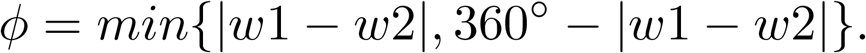

The corresponding absolute orientation difference *ω* of their strongest amplitude is equal to *ϕ*, for *ϕ* equal or less than 90^°^ otherwise, it is equal to 180^°^ − *ϕ*.

### Footnotes

[1 V1 receives sensory inputs in layer 4 (L4) processed vertically through the cortical column and laterally within each layer and then projected “forward” to higher areas by V1 layers 2/3 (L2/3), and “backwards” as feedback to lower areas by V1 L5/6.

[1 This is due to the discretization of the event rate per segment, which results from the specific number of calcium events that can occur during each segment.

[1 Due to space constraints, we do not present the linear relationships between the mutual information of a neuron with the stimulus and its z-scored MI (zMI), and FVE difference.

[1 This is a conservative threshold. A sensitivity analysis for lower thresholds is part of our ongoing research.

